# Presto scales Wilcoxon and auROC analyses to millions of observations

**DOI:** 10.1101/653253

**Authors:** Ilya Korsunsky, Aparna Nathan, Nghia Millard, Soumya Raychaudhuri

## Abstract

**Summary:** The related Wilcoxon rank sum test and area under the receiver operator curve are ubiquitous in high dimensional biological data analysis. Current implementations do not scale readily to the increasingly large datasets generated by novel high-throughput technologies, such as single cell RNAseq. We introduce a simple and scalable implementation of both analyses, available through the R package Presto. Presto scales to big datasets, with functions optimized for both dense and sparse matrices. On a sparse dataset of 1 million observations, 10 groups, and 1,000 features, Presto performed both rank-sum and auROC analyses in only 17 seconds, compared to 6.4 hours with base R functions. Presto also includes functions to seamlessly integrate with the Seurat single cell analysis pipeline and the Bioconductor SingleCellExperiment class. Presto enables the use of robust classical analyses on big data with a simple interface and optimized implementation.

**Availability and Implementation:** Presto is available as an R package at https://github.com/immunogenomics/presto.

**Supplementary Information:** Vignettes are available with the Presto package.

---

The Wilcoxon rank sum test (Wilcoxon, 1945; Pratt, 1959), also known as the Mann-Whitney U test, is a robust, nonparametric difference-of-means test. Since it assumes no underlying distribution, it is used widely in all fields of biology. Mathematically, it is closely related to a commonly-used metric, the area under the receiver operator curve (auROC). auROC is used to evaluate the power of a feature to distinguish two groups (Green and Swets, 1966). Both rank-sum and auROC are staple tools in high-dimensional genomics analyses, used for differential expression analysis in RNAseq and differential accessibility analysis in ATACseq. However, the current implementation of the rank-sum test in the wilcox.test function (R Core Team, 2019) cannot scale to the large datasets generated by single cell genomics technologies. Single cell RNAseq and single cell ATACseq datasets often contain data on millions of cells and thousands of features. At the same time, these large datasets often contain many zeros, with >90% sparsity in scRNAseq and >98% sparsity in scATACseq. Here, we present Presto, a fast implementation of the rank-sum test, auROC, and related summary statistics. Presto scales to millions of observations in seconds and has built-in optimizations for both dense and sparse datasets.

## Implementation details

We made use of several programmatic and mathematical tools to make Presto computationally efficient. We started with the vectorized multi-group approach to rank-sum in the MUDAN (Fan, 2018) and Pagoda2 (Kharchenko et al, 2019) R packages. In the multi-group setting, observations belong to 1 of K groups. We repeat rank-sum in a 1-versus-all fashion for each group individually. The vectorized implementation caches intermediate results that can be shared by each group. For instance, we rank the expression of a gene once across all groups and use this ranking to test each group individually. We implemented all computationally-intensive steps in C++. In these functions, we were careful to minimize the amount of memory allocation, performing many computations in place. Moreover, we wrote separate functions optimized for dense and sparse matrix inputs. For dense matrices, we used utilities available through the RcppArmadillo bindings (Eddelbuettel and Sanderson, 2014) to the C++ Armadillo package (Curtin and Sanderson, 2016; Curtin and Sanderson 2018). These functions are efficient for dense inputs, since Armadillo is built against a BLAS backend that enables fast dense matrix operations. While Armadillo has support for sparse matrices, this support is does not benefit from BLAS utilities. Therefore, we performed matrix operations directly on the “x”, “i”, and “p” fields defined in the dgCMatrix format, defined in the R Matrix package (Bates and Maechler, 2018). Finally, we exploited the simple mathematical connection between the rank-sum U statistic and auROC to compute auROC. For two groups of sizes N_1_ and N_2_, auROC = U / (N_1_*N_2_). Thus, once we compute the rank-sum U statistic, auROC is essentially computationally free.

We handled ties and computed p-values in the same way that the base R wilcox.test function does when run with parameters unpaired=TRUE, correct=TRUE, exact=FALSE, alternative=“two.sided”. The main front end function, wilcoxauc, is implemented in R. We implemented wilcoxauc using the S3 function-oriented class system. Here, wilcoxauc is a base function class with instantiations for Seurat objects, SingleCellExperiment objects, and matrix-like objects. For simplicity, we cast all dense matrix-like inputs (e.g. matrix, dgeMatrix, data.frame) as the base R matrix type and sparse inputs (e.g. dgCMatrix, dgTMatrix, TsparseMatrix) as the dgCMatrix type. The next section details how to use the wilcoxauc function in R.

## Usage

The Presto R package has a simple interface. Given a feature-by-observation matrix X and label vector y, Presto does a one-versus-all analysis for each group and each feature individually and reports a summary table of all tests.

library(devtools)

install_github(‘immunogenomics/presto’)

library(presto)

res <- wilcoxauc(X, y)

Presto also supports S4 objects commonly used in single cell genomics analysis, including Seurat (Stuart and Butler, 2018) and SingleCellExperiment (Lun and Risso, 2019). We assume these structures already have the covariates (e.g. gene expression matrix) and group labels (e.g. cluster IDs) defined internally. Presto has a common interface for S4 objects. The first argument is the object and the second the name of the group-defining variable.

res <- wilcoxauc(object, “label”)

For convenience, we include a utility function that summarizes the top abundant marker features for each group.

top_markers(res)

Finally, we note that many modern datasets, such as those generated with droplet based single cell RNAseq technologies (Macosko et al, 2015; Zheng et al, 2017), have a larger number of zero entries. These data can be compactly represented with a sparse matrix format, such as dgCMatrix, provided by the R Matrix package. Presto is optimized for both dense and sparse formats. When more than 30% of entries are zero, we recommend using the dgCMatrix format. More examples can be found in the R package documentation (?presto) and vignette.

## Benchmarking

We benchmarked the computational performance of Presto on an Intel Xeon E5-2690 v3 processor, with no parallelization. We compared the runtime of three implementations on simulated datasets of 10 groups, 1,000 features, and observations ranging from 100,000 (100K) to 1 million (1M). To demonstrate Presto’s performance with sparse data, we set 90% of entries to zero. For the base R (blue) implementation, we recorded runtimes from 38 minutes on 100K observations to 6.4 hours on 1 million. Presto analyzed the dense matrices in 7 to 83 seconds, with an average 295X speedup over base R. Presto analyzed the sparse matrices in only 2 to 16 seconds, with an average 1448X speedup over base R.

**Figure 1.**
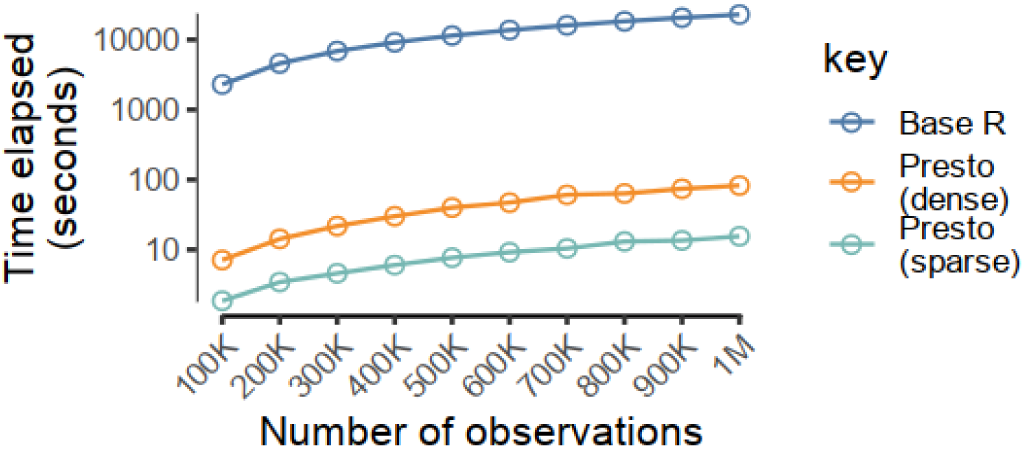
Runtime benchmarks using base R wilcox.test (blue), Presto dense (orange), and Presto sparse (teal). Datasets ranged from 100,000 to 1 million observations, with 1,000 features, 10 groups, and 90% sparsity. On 1 million cells, Presto on the sparse matrix finishes fastest, in only 16 seconds, compared to 84 seconds with dense Presto and 6.4 hours with base R.

## Conclusion

In this note, we present a modern implementation of classical methods. In the future, we plan to augment Presto with scalable versions of additional common tools. We hope this will allow analysts to scale their existing pipelines to large datasets with the minimum number of unnecessary technical interruptions.

## Supporting information

Vignette

## Acknowledgements

We thanks Sergio Poli and Tiffany Amariuta for beta testing Presto.

## Funding

This work is supported by funding from the National Institutes of Health (NIAMS T32AR007530, UH2AR067677, 1U01HG009088, U01HG009379, and 1R01AR063759).

## Notes

https://github.com/immunogenomics/presto

