## Supplementary material for "Presto scales Wilcoxon and auROC analyses to millions of observations": Vignette

Quick start to presto


Code 

- Show All Code
- Hide All Code

### Quick start to presto

Ilya Korsunsky

###### 28 May 2019

### Contents

- 1 Introduction
- 2 Installation
- 3 Input Types
  - 3.1 Matrix input
  - 3.2 Seurat
  - 3.3 SingleCellExperiment
- 4 Description of outputs
  - 4.1 Results table
  - 4.2 Top markers
- 5 Options
  - 5.1 Dense vs sparse
  - 5.2 groups\_use

### 1 Introduction

presto makes it fast and easy to run Wilcoxon rank sum test and auROC analysis
on large datasets. The tutorial below shows how to install presto, walks
through the 3 major ways you can use presto with your data, and finally
explores more advanced use cases.

### 2 Installation

To install the current stable release from CRAN:

```
install.packages('presto')
```

For the cutting edge version of presto:

```
library(devtools)
install_github('immunogenomics/presto')
```

### 3 Input Types

The main function in this vignette is `wilcoxauc`. presto currently supports 3
interfaces to `wilcoxauc`, with a matrix, Seurat object, or
SingleCellExperiment object. The output of `wilcoxauc` is described in the
next section.

```
?wilcoxauc
```

#### 3.1 Matrix input

The most general use of presto is with a matrix of features and observations
(exprs) paired with a vector of group labels (y).

```
data(exprs)
head(exprs)[, 1:10]
```

```
##    X1 X2 X3 X4 X5 X6 X7 X8 X9 X10
## G1  0  1  1  2  1  1  1  2  0   1
## G2  1  0  2  2  0  1  1  1  0   0
## G3  1  1  1  1  0  0  0  2  1   0
## G4  2  2  0  2  4  0  1  1  0   1
## G5  0  0  0  3  1  1  1  2  0   2
## G6  2  1  0  1  0  1  0  0  1   0
```

```
data(y)
head(y)
```

```
## [1] A A A A A A
## Levels: A B C
```

```
head(wilcoxauc(exprs, y))
```

```
##   feature group avgExpr  logFC statistic  auc  pval padj pct_in pct_out
## 1      G1     A    1.15  0.082      2890 0.55 0.257 0.64     75      59
## 2      G2     A    0.76 -0.300      2190 0.42 0.081 0.41     49      65
## 3      G3     A    0.98 -0.187      2429 0.46 0.452 0.75     65      68
## 4      G4     A    1.22  0.397      3176 0.61 0.020 0.41     71      56
## 5      G5     A    1.42  0.366      3026 0.58 0.093 0.41     73      57
## 6      G6     A    0.93 -0.178      2424 0.46 0.439 0.75     56      60
```

#### 3.2 Seurat

We support interfacing with Seurat Version 3 objects. In the most basic use
case, specify the Seurat object and the meta data variable that defines the
group labels.

```
data(object_seurat)
head(wilcoxauc(object_seurat, 'cell_type'))
```

```
##   feature  group avgExpr   logFC statistic  auc pval padj pct_in pct_out
## 1      G1 jurkat     6.0  0.0406     11343 0.50 0.90 1.00    100     100
## 2      G2 jurkat     5.9 -0.0278     11209 0.50 0.96 1.00    100     100
## 3      G3 jurkat     5.9 -0.0584     10306 0.46 0.21 0.99    100     100
## 4      G4 jurkat     5.9 -0.0597     10859 0.48 0.61 1.00    100     100
## 5      G5 jurkat     6.0 -0.0202     11244 0.50 1.00 1.00    100     100
## 6      G6 jurkat     5.9 -0.0075     11038 0.49 0.78 1.00    100     100
```

Seurat supports multiple assay types. This can be specified with
`seurat_assay`.

```
head(wilcoxauc(object_seurat, 'cell_type', seurat_assay = 'RNA'))
```

```
##   feature  group avgExpr   logFC statistic  auc pval padj pct_in pct_out
## 1      G1 jurkat     6.0  0.0406     11343 0.50 0.90 1.00    100     100
## 2      G2 jurkat     5.9 -0.0278     11209 0.50 0.96 1.00    100     100
## 3      G3 jurkat     5.9 -0.0584     10306 0.46 0.21 0.99    100     100
## 4      G4 jurkat     5.9 -0.0597     10859 0.48 0.61 1.00    100     100
## 5      G5 jurkat     6.0 -0.0202     11244 0.50 1.00 1.00    100     100
## 6      G6 jurkat     5.9 -0.0075     11038 0.49 0.78 1.00    100     100
```

Seurat supports multiple feature expression matrices, such as raw counts,
library normalized data, and scaled data. These can be accessed with `assay`.

```
head(wilcoxauc(object_seurat, 'cell_type', assay = 'counts'))
```

```
##   feature  group avgExpr   logFC statistic  auc pval padj pct_in pct_out
## 1      G1 jurkat    0.51  0.0046     11385 0.51 0.85 0.97    100     100
## 2      G2 jurkat    0.52 -0.0032     11221 0.50 0.97 0.97    100     100
## 3      G3 jurkat    0.48 -0.0373     10398 0.46 0.26 0.97    100     100
## 4      G4 jurkat    0.51 -0.0172     10865 0.48 0.61 0.97    100     100
## 5      G5 jurkat    0.51  0.0015     11301 0.50 0.94 0.97    100     100
## 6      G6 jurkat    0.50 -0.0065     11137 0.50 0.89 0.97    100     100
```

```
head(wilcoxauc(object_seurat, 'cell_type', assay = 'data'))
```

```
##   feature  group avgExpr   logFC statistic  auc pval padj pct_in pct_out
## 1      G1 jurkat     6.0  0.0406     11343 0.50 0.90 1.00    100     100
## 2      G2 jurkat     5.9 -0.0278     11209 0.50 0.96 1.00    100     100
## 3      G3 jurkat     5.9 -0.0584     10306 0.46 0.21 0.99    100     100
## 4      G4 jurkat     5.9 -0.0597     10859 0.48 0.61 1.00    100     100
## 5      G5 jurkat     6.0 -0.0202     11244 0.50 1.00 1.00    100     100
## 6      G6 jurkat     5.9 -0.0075     11038 0.49 0.78 1.00    100     100
```

```
head(wilcoxauc(object_seurat, 'cell_type', assay = 'scale.data'))
```

```
##   feature  group avgExpr   logFC statistic  auc pval padj pct_in pct_out
## 1      G1 jurkat  0.0206  0.0420     11343 0.50 0.90 1.00    100     100
## 2      G2 jurkat -0.0140 -0.0286     11209 0.50 0.96 1.00    100     100
## 3      G3 jurkat -0.0329 -0.0671     10306 0.46 0.21 0.99    100     100
## 4      G4 jurkat -0.0314 -0.0641     10859 0.48 0.61 1.00    100     100
## 5      G5 jurkat -0.0114 -0.0232     11244 0.50 1.00 1.00    100     100
## 6      G6 jurkat -0.0038 -0.0077     11038 0.49 0.78 1.00    100     100
```

#### 3.3 SingleCellExperiment

presto supports the Bioconductor data structure SingleCellExperiment. Again,
the most simple use case takes a SingleCellExperiment object and the metadata
field with group labels.

```
data(object_sce)
head(wilcoxauc(object_sce, 'cell_type'))
```

```
## Loading required package: SingleCellExperiment
```

```
## Loading required package: SummarizedExperiment
```

```
## Loading required package: GenomicRanges
```

```
## Loading required package: stats4
```

```
## Loading required package: BiocGenerics
```

```
## Loading required package: parallel
```

```
## 
## Attaching package: 'BiocGenerics'
```

```
## The following objects are masked from 'package:parallel':
## 
##     clusterApply, clusterApplyLB, clusterCall, clusterEvalQ,
##     clusterExport, clusterMap, parApply, parCapply, parLapply,
##     parLapplyLB, parRapply, parSapply, parSapplyLB
```

```
## The following objects are masked from 'package:stats':
## 
##     IQR, mad, sd, var, xtabs
```

```
## The following objects are masked from 'package:base':
## 
##     anyDuplicated, append, as.data.frame, basename, cbind, colMeans,
##     colnames, colSums, dirname, do.call, duplicated, eval, evalq,
##     Filter, Find, get, grep, grepl, intersect, is.unsorted, lapply,
##     lengths, Map, mapply, match, mget, order, paste, pmax, pmax.int,
##     pmin, pmin.int, Position, rank, rbind, Reduce, rowMeans,
##     rownames, rowSums, sapply, setdiff, sort, table, tapply, union,
##     unique, unsplit, which, which.max, which.min
```

```
## Loading required package: S4Vectors
```

```
## 
## Attaching package: 'S4Vectors'
```

```
## The following object is masked from 'package:base':
## 
##     expand.grid
```

```
## Loading required package: IRanges
```

```
## Loading required package: GenomeInfoDb
```

```
## Loading required package: Biobase
```

```
## Welcome to Bioconductor
## 
##     Vignettes contain introductory material; view with
##     'browseVignettes()'. To cite Bioconductor, see
##     'citation("Biobase")', and for packages 'citation("pkgname")'.
```

```
## Loading required package: DelayedArray
```

```
## Loading required package: matrixStats
```

```
## 
## Attaching package: 'matrixStats'
```

```
## The following objects are masked from 'package:Biobase':
## 
##     anyMissing, rowMedians
```

```
## Loading required package: BiocParallel
```

```
## 
## Attaching package: 'DelayedArray'
```

```
## The following objects are masked from 'package:matrixStats':
## 
##     colMaxs, colMins, colRanges, rowMaxs, rowMins, rowRanges
```

```
## The following objects are masked from 'package:base':
## 
##     aperm, apply
```

```
##   feature  group avgExpr   logFC statistic  auc pval padj pct_in pct_out
## 1      G1 jurkat     6.0  0.0406     11343 0.50 0.90 1.00    100     100
## 2      G2 jurkat     5.9 -0.0278     11209 0.50 0.96 1.00    100     100
## 3      G3 jurkat     5.9 -0.0584     10306 0.46 0.21 0.99    100     100
## 4      G4 jurkat     5.9 -0.0597     10859 0.48 0.61 1.00    100     100
## 5      G5 jurkat     6.0 -0.0202     11244 0.50 1.00 1.00    100     100
## 6      G6 jurkat     5.9 -0.0075     11038 0.49 0.78 1.00    100     100
```

SingleCellExperiment can have several data slots, such as counts and logcounts.
These can be accessed with `assay`.

```
head(wilcoxauc(object_sce, 'cell_type', assay = 'counts'))
```

```
##   feature  group avgExpr   logFC statistic  auc pval padj pct_in pct_out
## 1      G1 jurkat    0.51  0.0046     11385 0.51 0.85 0.97    100     100
## 2      G2 jurkat    0.52 -0.0032     11221 0.50 0.97 0.97    100     100
## 3      G3 jurkat    0.48 -0.0373     10398 0.46 0.26 0.97    100     100
## 4      G4 jurkat    0.51 -0.0172     10865 0.48 0.61 0.97    100     100
## 5      G5 jurkat    0.51  0.0015     11301 0.50 0.94 0.97    100     100
## 6      G6 jurkat    0.50 -0.0065     11137 0.50 0.89 0.97    100     100
```

```
head(wilcoxauc(object_sce, 'cell_type', assay = 'logcounts'))
```

```
##   feature  group avgExpr   logFC statistic  auc pval padj pct_in pct_out
## 1      G1 jurkat     6.0  0.0406     11343 0.50 0.90 1.00    100     100
## 2      G2 jurkat     5.9 -0.0278     11209 0.50 0.96 1.00    100     100
## 3      G3 jurkat     5.9 -0.0584     10306 0.46 0.21 0.99    100     100
## 4      G4 jurkat     5.9 -0.0597     10859 0.48 0.61 1.00    100     100
## 5      G5 jurkat     6.0 -0.0202     11244 0.50 1.00 1.00    100     100
## 6      G6 jurkat     5.9 -0.0075     11038 0.49 0.78 1.00    100     100
```

### 4 Description of outputs

#### 4.1 Results table

All inputs for `wilcoxauc` give the same table of results.

| parameter | description |
| --- | --- |
| feature | name of feature. |
| group | name of group label. |
| avgExpr | mean value of feature in group. |
| logFC | log fold change between observations in group vs out. |
| statistic | Wilcoxon rank sum U statistic. |
| auc | area under the receiver operator curve. |
| pval | nominal p value, from two-tailed Gaussian approximation of U statistic. |
| padj | Benjamini-Hochberg adjusted p value. |
| pct\_in | Percent of observations in the group with non-zero feature value. |
| pct\_out | Percent of observations out of the group with non-zero feature value. |

```
head(wilcoxauc(exprs, y))
```

```
##   feature group avgExpr  logFC statistic  auc  pval padj pct_in pct_out
## 1      G1     A    1.15  0.082      2890 0.55 0.257 0.64     75      59
## 2      G2     A    0.76 -0.300      2190 0.42 0.081 0.41     49      65
## 3      G3     A    0.98 -0.187      2429 0.46 0.452 0.75     65      68
## 4      G4     A    1.22  0.397      3176 0.61 0.020 0.41     71      56
## 5      G5     A    1.42  0.366      3026 0.58 0.093 0.41     73      57
## 6      G6     A    0.93 -0.178      2424 0.46 0.439 0.75     56      60
```

#### 4.2 Top markers

We often find it helpful to summarize what the most distinguishing features
are in each group.

```
res <- wilcoxauc(exprs, y)
top_markers(res, n = 10)
```

```
## # A tibble: 10 x 4
##     rank A     B     C    
##    <int> <chr> <chr> <chr>
##  1     1 G4    G20   G1   
##  2     2 G5    G5    G21  
##  3     3 G9    G15   G6   
##  4     4 G10   G19   G16  
##  5     5 G14   G25   G11  
##  6     6 G1    G10   G7   
##  7     7 G15   G13   G2   
##  8     8 G25   G8    G17  
##  9     9 G8    G24   G12  
## 10    10 G24   G23   G3
```

We can also filter for some criteria. For instance, the top features must be
in at least 70% of all observations within the group. Note that not all groups
have 10 markers that meet these criteria.

```
res <- wilcoxauc(exprs, y)
top_markers(res, n = 10, auc_min = .5, pct_in_min = 70)
```

```
## # A tibble: 9 x 4
##    rank A     B     C    
##   <int> <chr> <chr> <chr>
## 1     1 G4    G20   G1   
## 2     2 G5    G5    G21  
## 3     3 G14   G15   G6   
## 4     4 G1    G19   G16  
## 5     5 <NA>  G25   G11  
## 6     6 <NA>  G23   G7   
## 7     7 <NA>  G9    G2   
## 8     8 <NA>  G14   G12  
## 9     9 <NA>  <NA>  G3
```

### 5 Options

#### 5.1 Dense vs sparse

presto is optimized for dense and sparse matrix inputs. When possible, use
sparse inputs. In our toy dataset, almost 39% of elements are zeros. Thus, it
makes sense to cast it as a sparse dgCMatrix and run wilcoxauc on that.

```
sum(exprs == 0) / prod(dim(exprs))
```

```
## [1] 0.39
```

```
exprs_sparse <- as(exprs, 'dgCMatrix')
head(wilcoxauc(exprs_sparse, y))
```

```
##   feature group avgExpr  logFC statistic  auc  pval padj pct_in pct_out
## 1      G1     A    1.15  0.082      2890 0.55 0.257 0.64     75      59
## 2      G2     A    0.76 -0.300      2190 0.42 0.081 0.41     49      65
## 3      G3     A    0.98 -0.187      2429 0.46 0.452 0.75     65      68
## 4      G4     A    1.22  0.397      3176 0.61 0.020 0.41     71      56
## 5      G5     A    1.42  0.366      3026 0.58 0.093 0.41     73      57
## 6      G6     A    0.93 -0.178      2424 0.46 0.439 0.75     56      60
```

#### 5.2 groups\_use

Sometimes, you don’t want to test all groups in the dataset against all other
groups. For instance, I want to compare only observations in group ‘A’ to
those in group ‘B’. This is achieved with the groups\_use argument.

```
res_AB <- wilcoxauc(exprs, y, groups_use = c('A', 'B'))
head(res_AB)
```

```
##   feature group avgExpr  logFC statistic  auc    pval    padj pct_in pct_out
## 1      G1     A    1.15  0.790      1868 0.75 2.4e-06 0.00006     75      29
## 2      G2     A    0.76 -0.081      1184 0.48 6.9e-01 0.79372     49      53
## 3      G3     A    0.98 -0.018      1246 0.50 9.5e-01 0.97606     65      62
## 4      G4     A    1.22  0.240      1413 0.57 2.0e-01 0.44123     71      60
## 5      G5     A    1.42 -0.248      1054 0.43 1.9e-01 0.44123     73      78
## 6      G6     A    0.93  0.438      1521 0.61 3.1e-02 0.14516     56      36
```

```
top_markers(res_AB)
```

```
## # A tibble: 10 x 3
##     rank A     B    
##    <int> <chr> <chr>
##  1     1 G1    G20  
##  2     2 G21   G19  
##  3     3 G16   G13  
##  4     4 G6    G15  
##  5     5 G11   G5   
##  6     6 G4    G25  
##  7     7 G12   G18  
##  8     8 G17   G23  
##  9     9 G22   G8   
## 10    10 G9    G10
```
